# Integrating mechanism-based T cell phenotypes into a model of tumor-immune cell interactions

**DOI:** 10.1101/2024.03.05.583613

**Authors:** Neel Tangella, Colin G. Cess, Geena V. Ildefonso, Stacey D. Finley

## Abstract

Interactions between cancer cells and immune cells in the tumor microenvironment influence tumor growth and can contribute to the response to cancer immunotherapies. It is difficult to gain mechanistic insights into the effects of cell-cell interactions in tumors using a purely experimental approach. However, computational modeling enables quantitative investigation of the tumor microenvironment, and agent-based modeling in particular provides relevant biological insights into the spatial and temporal evolution of tumors. Here, we develop a novel agent-based model (ABM) to predict the consequences of intercellular interactions. Furthermore, we leverage our prior work that predicts the transitions of CD8+ T cells from a naïve state to a terminally differentiated state using Boolean modeling. Given the detailed incorporated to predict T cell state, we apply the integrated Boolean-ABM framework to study how the properties of CD8+ T cells influence the composition and spatial organization of tumors and the efficacy of an immune checkpoint blockade. Overall, we present a mechanistic understanding of tumor evolution that can be leveraged to study targeted immunotherapeutic strategies.

## 1 INTRODUCTION

The tumor microenvironment (TME) is a complex ecosystem comprised of many different cell types, including cancer cells and immune cells. It has long been shown that the immune system plays a pivotal role in tumor development.^1–4^ Although immune cells can detect malignant cells as “not-self” and then eliminate them, tumors are still able to grow and escape the immune system. In fact, in several cancer types, the TME has been shown to strongly contribute to the immunosuppressive state of the tumor. Furthermore, several studies point to the importance of the spatial location of stromal and immune cells in predicting prognosis and treatment response.^5,6^ Thus, by considering the composition and spatial structure of tumors, it may be possible to better understand the factors that drive tumor progression and the efficacy of anti-cancer treatments.^7^

Given its complexity, efforts to extensively explore the TME using *in vitro* and *in vivo* models alone are intractable. Fortunately, computational modeling is a useful approach to study the TME and understand how individual cell-cell interactions and changes at the molecular and cellular levels influence tumor growth and response to treatment.^8^ For example, computational models have been used to better understand the interactions within the TME and predict how to improve immune-based cancer therapies.^9^ A range of computational models have been developed to study various aspects of the TME and immunotherapy.^10–13^ Agent-based models (ABMs) have proven to be particularly useful,^14–16^ as they predict the spatial and temporal evolution of tumors and can simulate tumor behavior for a range of different conditions that are difficult and time-consuming to study using a purely experimental approach.

While many ABMs exist to study the TME, the large majority use phenomenological rules to determine the cells’ behaviors.^15,16^ Such a rule-based approach does not explicitly account for the intracellular mechanisms (i.e., signaling transduction, metabolic pathways, transcriptional regulation) that inform a cell’s decision to proliferate, change phenotypic states, or induce cell killing. Here, we build on our prior work using biophysical agent-based modeling to simulate tumor growth^17–19^ by integrating our published Boolean model^20^ of a gene regulatory network (GRN) shown to mediate T cell exhaustion. We predict the evolution of phenotypic T cell states, informed by the GRN encoded in the Boolean model. We consider three T cell states (naïve, pro-memory, and exhausted), which each have distinct properties. We apply the integrated model to study how CD8+ T cell state and properties influence tumor composition and spatial organization. Furthermore, we simulate immunotherapy via blockade of programmed cell death protein 1 (PD1), which is actively being pursued as a target to control tumor growth.^21,22^ By accounting for the GRN governing T cell behavior, which influences cell-cell interactions, we provide a mechanistic view of tumor evolution and provide a framework for predicting how tumor growth can be controlled via genetic alterations of T cells.

## 2 RESULTS

Here, we use a novel model of tumor growth that integrates a the GRN of CD8+ T cell states (predicted by a Boolean model) into an ABM that predicts cell-level behaviors and cell-cell interactions. This model leverages our prior work in simulating T cell state transitions.^20^ Additionally, the ABM represents a significant expansion of our published modeling of cell-cell interactions within the tumor microenvironment.^17–19^ We apply the novel modeling framework to simulate tumor growth, with varying CD8+ T cell properties.

### The rate of CD8+ T cell recruitment alters the number of cancer cells and relative proportions of immune cell subsets

We utilized this model and considered the baseline values for the probabilities of CD8+ T cell death and cancer cell killing and investigated the effects of the rate of CD8+ T cell recruitment. Overall, the model predicts tumor escape and the presence of pro-tumor immune cells for these CD8+ T cell properties. Panel A of **Figures 1 and 2** shows the number of cancer cells over time averaged over 10 simulation replicates, as well as the standard deviation. The model predicts that higher CD8+ T cell recruitment leads to fewer cancer cells (30,124 and 22,357 for low and high recruitment rates, respectively, at the end of the model simulation).

**Figure 1.**
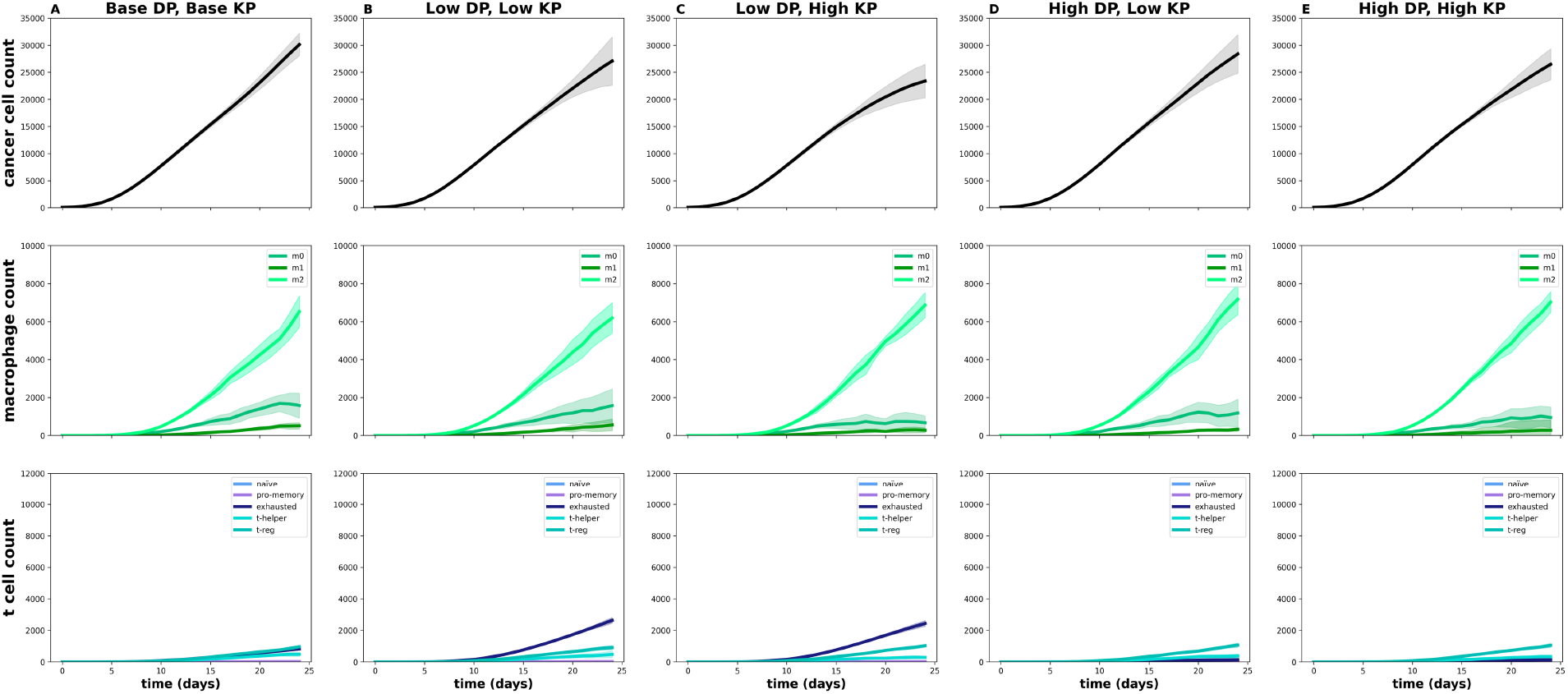
Time courses of cell counts for various CD8+ T cell properties, with base CD8+ T cell recruitment rate. *dp*, probability of CD8+ T cell death; *kp*, probability of CD8+ T cell-mediated cancer cell killing. Note the *y*-axes limits are set to enable direct comparison with Figures 2, 7, and S2.

**Figure 2.**
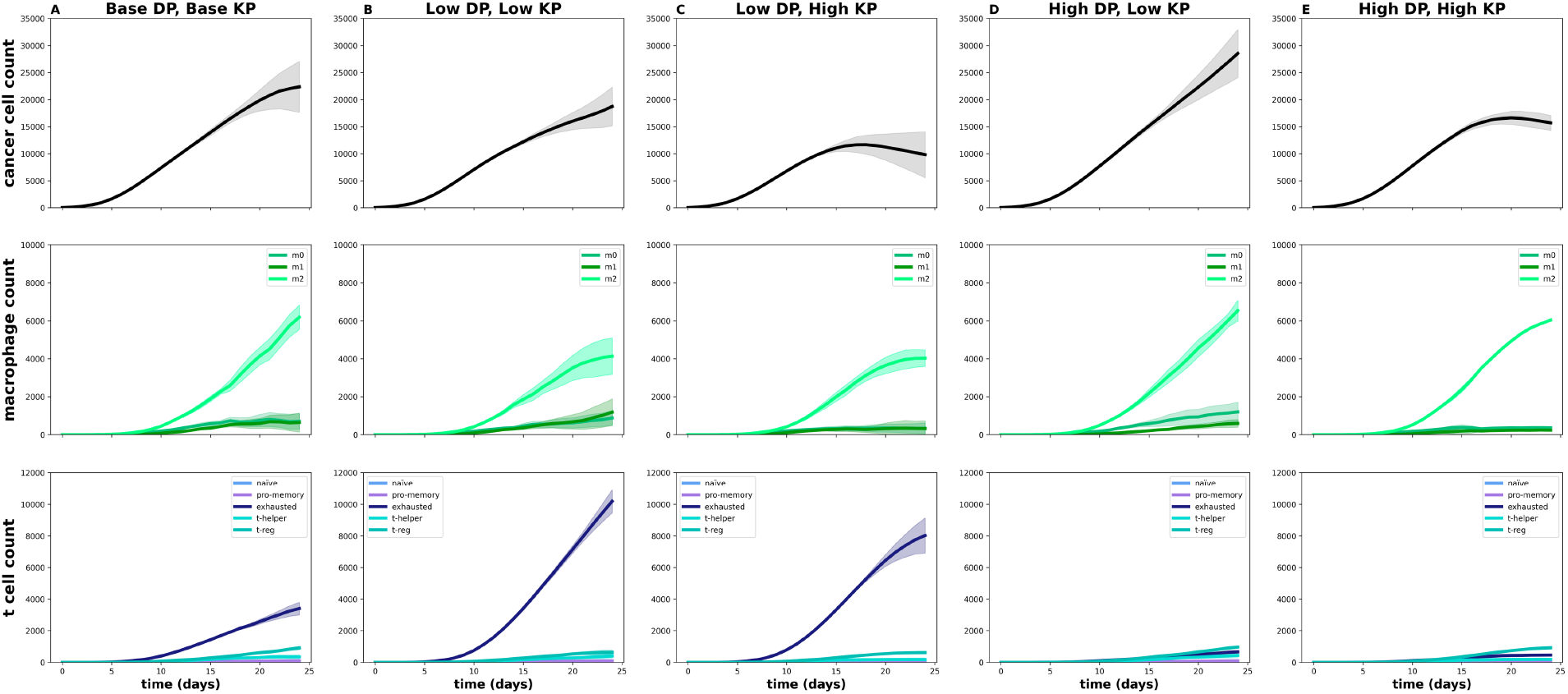
Time courses of cell counts for various CD8+ T cell properties, with high CD8+ T cell recruitment rate. *dp*, probability of CD8+ T cell death; *kp*, probability of CD8+ T cell-mediated cancer cell killing. Note the *y*-axes limits are set to enable direct comparison with Figures 1, 7, and S2.

The final numbers of macrophages are predicted to be slightly higher for high CD8+ T cell recruitment (8,603 and 7,525 macrophages for low and high recruitment rate, respectively). Similarly, the final relative proportion of macrophage types is only moderately impacted by the CD8+ T cell recruitment rate (**Figure 3**). Specifically, pro-tumor macrophages comprise 76% of the macrophage population for low T cell recruitment, compared to 82% for fast T cell recruitment.

**Figure 3.**
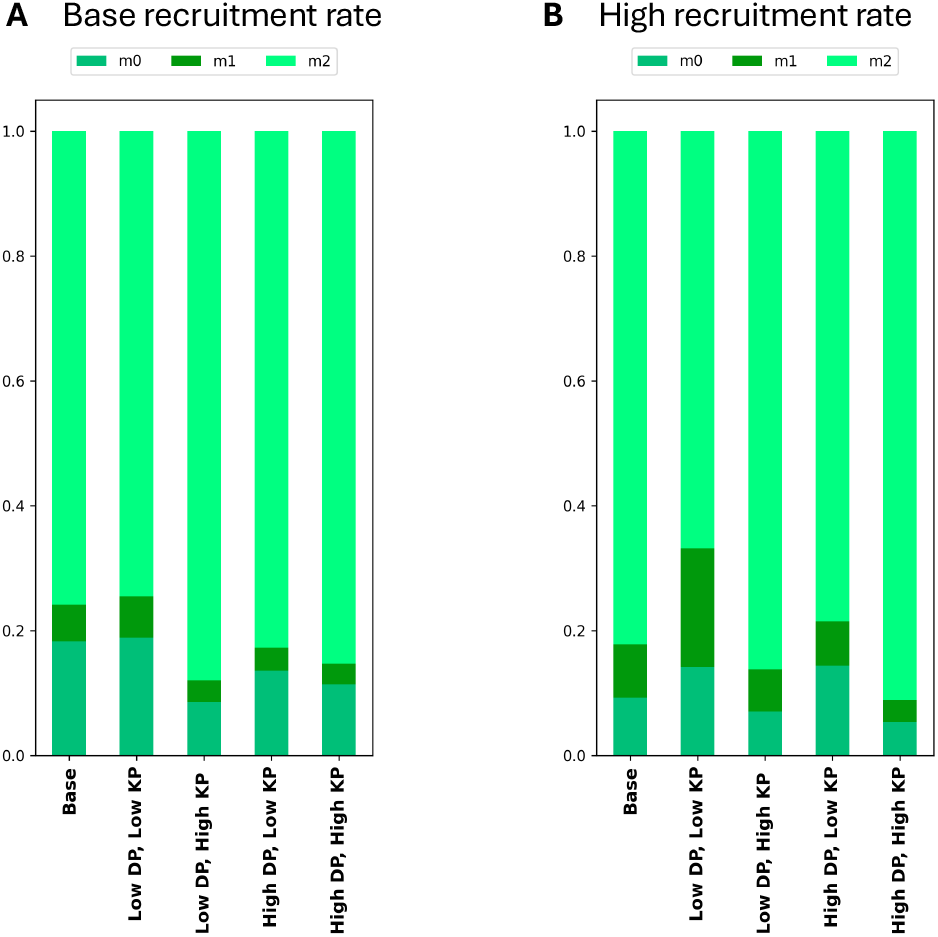
Relative proportions of macrophages at the end of the model simulation for various CD8+ T cell properties. **A**, Base CD8+ T cell recruitment rate. **B**, High CD8+ T cell recruitment rate. *dp*, probability of CD8+ T cell death; *kp*, probability of CD8+ T cell-mediated cancer cell killing.

Furthermore, the number of exhausted CD8+ T cells increases by nearly four-fold (from 810to 3,396 for low and high T cell recruitment, respectively). This corresponds to differences in the relative proportions of T cell subsets: 36% of CD8+ T cells are in an exhausted state for a low recruitment rate, compared to 72% for a high recruitment rate (**Figure 4**). In addition, CD4+ T helper cells comprise 21% of the T cell compartment for low T cell recruitment, but only 7% for high T cell recruitment. These changes in the subsets of immune cells make sense, as faster CD8+ T cell recruitment enables more T cells to enter the tumor; however, those T rapidly evolve towards the exhausted state. A larger proportion of exhausted CD8+ T cells corresponds to fewer CD4+ helper cells, leading to fewer anti-tumor (M1) macrophages, due to T cell-macrophage interactions. We note that across both CD8+ T cell recruitment rates, naïve and pro-memory T cells represent less than 3% of the T cell compartment.

**Figure 4.**
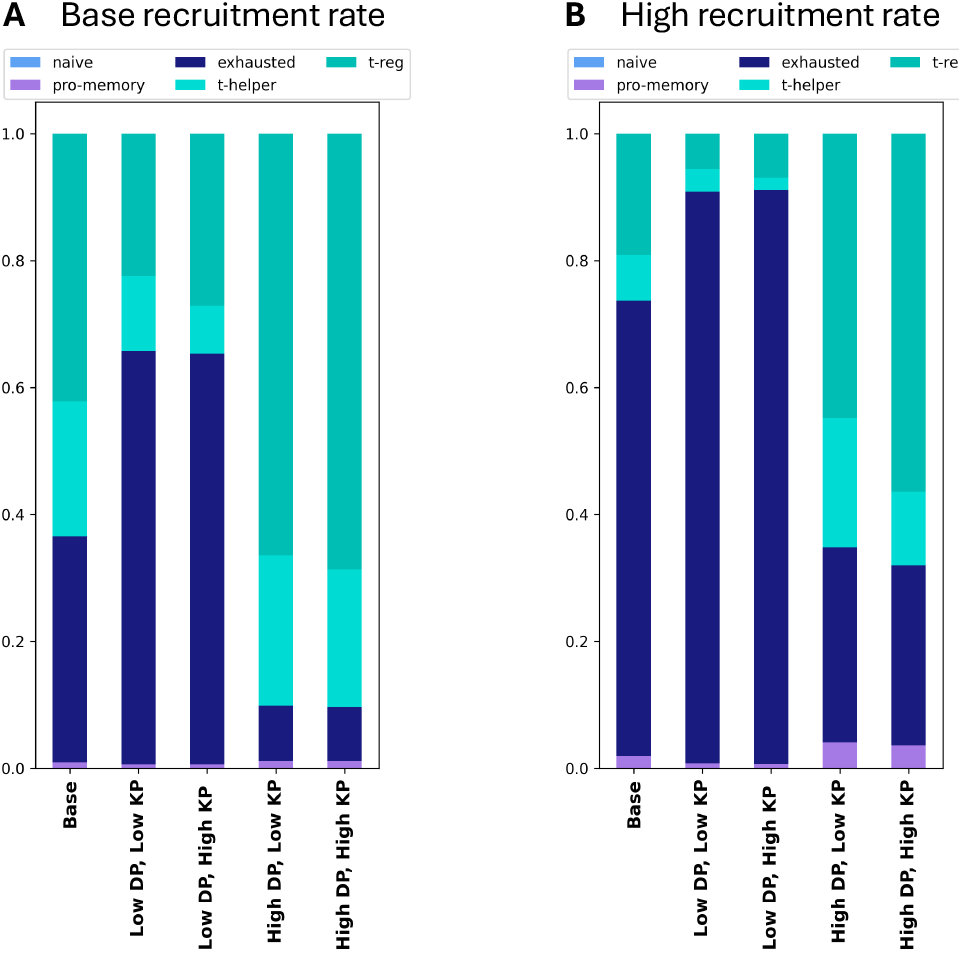
Relative proportions of T cells at the the end of the model simulation for various CD8+ T cell properties. **A**, Base CD8+ T cell recruitment rate. **B**, High CD8+ T cell recruitment rate. *dp*, probability of CD8+ T cell death; *kp*, probability of CD8+ T cell-mediated cancer cell killing.

A useful feature of ABMs is their ability to predict the spatial organization of the cell populations being modeled. Thus, in addition to considering the time courses of the cell populations, we also examined the spatial layout at the end of the time course (**Figure 5**) and over time (Supplementary Materials, Movies 1 and 2). The predictions show that for both CD8+ T cell recruitment rates considered, there is minimal infiltration of CD8+ T cells, as they remain at the periphery of the tumor. This aligns with the concept of most tumors being classified as immunologically “cold”, defined as the having low T cell infiltration.^23^ The tumor periphery is also rich in macrophages. Additionally, there is moderate infiltration of macrophages into the tumor, and there is slightly more macrophage infiltration for the case of low CD8+ T cell recruitment rate. The model predicts that the average distance of macrophages from the tumor center is 1930 *μ*m and 1647 *μ*m for low and high recruitment rate, respectively (**Table S1**).

**Figure 5.**
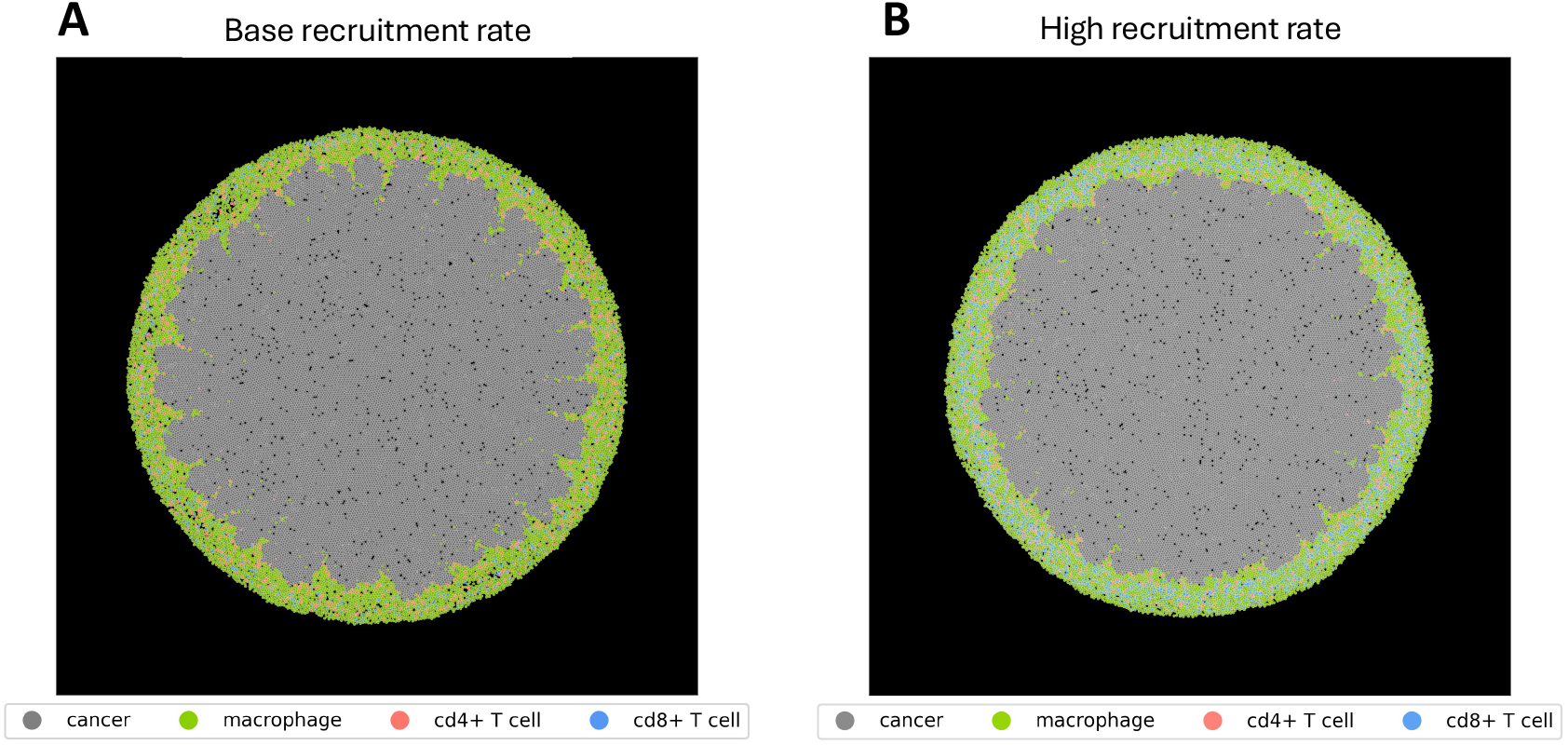
Representative tumor spatial layouts for baseline model at the end of the simulation. **A**, Base CD8+ T cell recruitment rate. **B**, High CD8+ T cell recruitment rate.

### When CD8+ T cell recruitment is low, T cell death and cytotoxicity have differential effects on the number of cancer cells and the proportion of immune cells

The model predicts that the size of the cancer cell population is largely insensitive to the CD8+ T cell properties investigated. Here, we return to the analysis of the time courses shown in **Figure 1**. The number of cancer cells increases exponentially for both low and high probability of CD8+ T cell death, as well as the two probabilities of T cell-mediated cancer cell killing (**Figure 1B-E**, top row). Furthermore, the number of cancer cells present at the end of the simulation is approximately the same across all conditions. Similarly, the spatial organization of the tumor is consistent across CD8+ T cell properties (Supplementary Materials, Figure S1).

In comparison, the number and relative proportions of macrophages depend on CD8+ T cell properties. The model predicts increases in the number of pro-tumor macrophages over time, for all four conditions (**Figure1B-E**, middle row). When CD8+ T cells have a low probability of cell death, M2 cells comprise 75-88% of the macrophage compartment (**Figure 3**). However, there are fewer M2 cells when the CD8+ T cells have a high probability of cell death, comprising 80-85% of macrophages, for high or low probability of CD8+ T cell-mediated cancer cell killing, respectively.

As anticipated, the probability of CD8+ T cell death and the probability of inducing cancer cell killing, influences the number of T cells (**Figure1B-E**, bottom row) and relative proportion of T cells in the tumor (**Figure 4**). When CD8+ T cells have a low probability of cell death, exhausted CD8+ T cells, Tregs, and CD4+ T helper cells make up approximately 65%, 25%, and 10% of the T cell population, respectively. In contrast, with high probability of CD8+ T cell death, the proportion of exhausted CD8+ T cell death is limited to 9% of all T cells. Having a smaller relative proportion of exhausted T cells corresponds to greater fractions of Tregs and T helper cells, which range from 66-69% and 22-24%, respectively, depending on the probability of T cell-mediated cancer cell killing. Again, these results can be traced to the cell-cell interactions included in the model. Having fewer pro-tumor macrophages (in the condition of high probability of T cell death), corresponds to more CD4+ T helper cells, which promotes differentiation of naïve macrophages into the anti-tumor state (M1).

Overall, the model predicts that with low recruitment of CD8+ T cells, varying the properties of the CD8+ T cells influences the number and proportions of immune cells, with only a minimal effect on the number of cancer cells. This is not an intuitive result, demonstrating the utility of a systems-level model of the tumor microenvironment.

### High CD8+ T cell recruitment permits tumor control and increases the relative size of the anti-tumor immune cell population

Across a range of values for the probability of CD8+ T cell death and cancer cell killing probability, the final number of cancer cells is lower for fast CD8+ T cell recruitment, compared to the baseline recruitment rate (**Figures 1 and 2**, top row). One exception is the condition of a high probability of CD8+ T cell death combined with a low probability of T cell-mediated killing (panel D in **Figures 1 and 2**), as compared to the baseline rate of CD8+ T cell recruitment.

The faster rate of CD8+ T cell recruitment is predicted to influence the absolute number and relative proportions of macrophage and T cell subsets in the tumor. For all four conditions, the number of pro-tumor macrophages is lower when T cell recruitment is high (**Figures 1 and 2**, middle row). Additionally, anti-tumor macrophages comprise a larger fraction of the macrophage population, (4-19% and 3-7% for high and base recruitment rates, respectively), as shown in **Figure 3**. Furthermore, increasing CD8+ T cell recruitment leads to a larger number (**Figures 1 and 2**, middle row) and a greater proportion (**Figure 4**) of exhausted CD8+ T cells across all conditions. For example, exhausted CD8+ T cells comprise 9% for the base T cell recruitment rate, compared to 28% for the case of high recruitment. These changes correspond to a smaller fraction of Tregs and greater percentage of pro-memory CD8+ T cells. Altogether, the model predicts that higher CD8+ T cell recruitment increases the proportion of anti-tumor immune cells (M1, pro-memory CD8+ T cells), concomitant with a decrease in the relative size of the anti-tumor immune cell population (M2 and Tregs).

The spatial layout of the tumor further demonstrates the effect of increased CD8+ T cell recruitment (**Figure 6**). A comparison of the spatial organization of the tumor for base and high recruitment rates for a representative model output shows fewer cancer cells, with the exception of the case for the high probability of CD8+ T cell death combined with low probability of T cell-mediated killing (**Figure 6C, G**). The tumors appear smaller, and we can visually appreciate the presence of substantially more exhausted CD8+ T cells (colored blue). Additionally, there is greater infiltration of immune cells into the tumor when CD8+ T cells are recruited at a faster rate and have a low probability of death (**Figure 6E, G** and **Table S1**).

**Figure 6.**
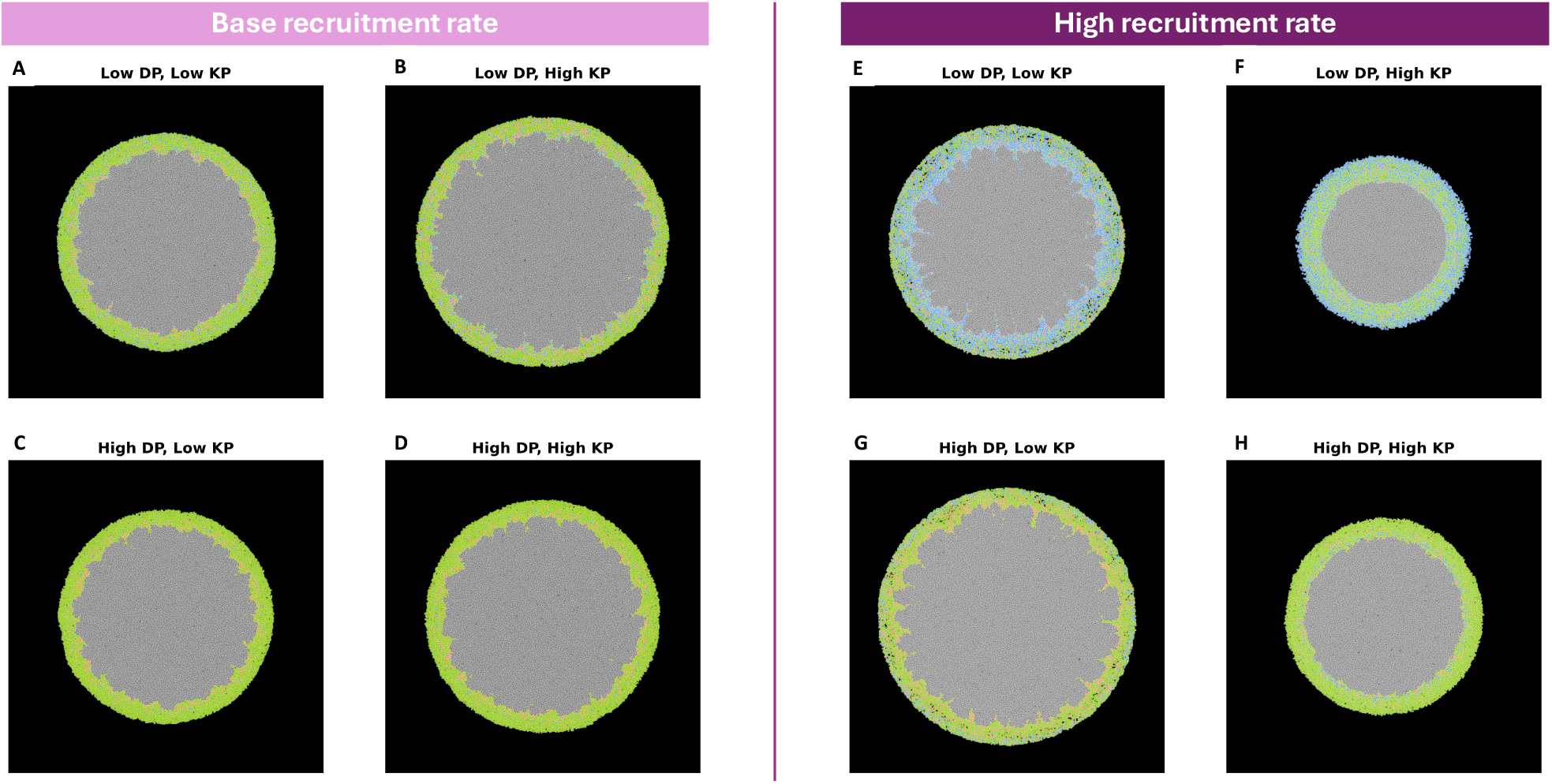
Representative tumor spatial layouts for various CD8+ T cell properties at the end of the simulation. **Left**, Base CD8+ T cell recruitment rate. **Right**, High CD8+ T cell recruitment rate. *dp*, probability of CD8+ T cell death; *kp*, probability of CD8+ T cell-mediated cancer cell killing.

### PD1 blockade increases the number of pro-memory CD8+ T cells and enables better tumor control when T cell recruitment is high

Finally, we considered the effects of altering the intracellular GRN of CD8+ T cells compared to the baseline model (termed wildtype, or “WT”). In particular, we simulated inhibition of PD1. Experimental studies demonstrate that blocking the PD1 pathway in naïve CD8+ T cells during differentiation can alter the cell state transitions. As in our previous work, we implement PD1 blockade by removing the ability of NFATC1 to activate the PD1 in the Boolean model of the CD8+ T cell GRN.

When CD8+ T cells are recruited to the tumor at the base rate, the model predicts that inhibiting PD1 strongly decreases the absolute number of exhausted CD8+ T cells, versus the WT case (compare the bottom rows of **Figures 1 and S1**). Additionally, PD1 blockade significantly increases the proportion of pro-memory CD8+ T cells (compare **Figure 4A** to **Figure S1B**). Despite altering the composition of the T cell population, when CD8+ T cell recruitment is low, PD1 blockade does not affect the number of tumor cells or macrophages (compare the top two rows in **Figures 1 and S2**). PD1 blockade also fails to alter the relative proportions of macrophages at the end of the model simulation (compare **Figure 3A** to **Figure S1A**).

In contrast, with higher CD8+ T cell recruitment, the model predicts that PD1 blockade can lead to smaller tumors compared to WT with high T cell recruitment. In particular, for the base and low values of the probability of CD8+ T cells considered, the numbers of tumor cells and pro-tumor macrophages are predicted to be lower than WT (compare the top two rows in **Figures 2 and 7**). For example, with the baseline values for the probability of CD8+ T cell death and T cell-mediated cancer cell killing, the number of cancer cells is 22,357 and 15,498 for WT and PD1 blockade, respectively. Interestingly, while the absolute numbers of M2 cells decrease with PD1 blockade, this does not always correspond to a change in the absolute number (compare bottom row of **Figures 2 and 7**) or final relative proportion of macrophages (compare **Figures 3B and 8A**).

**Figure 7.**
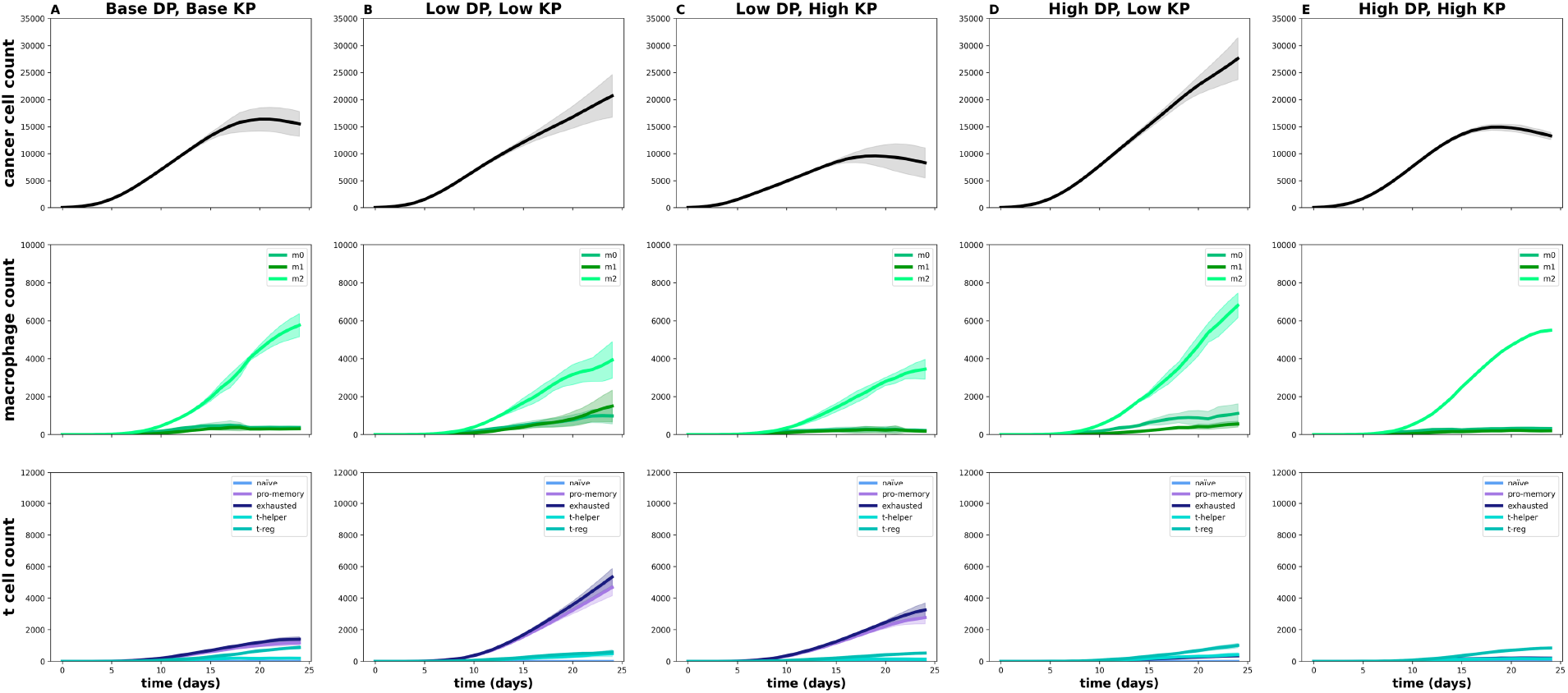
Time courses of cell counts for various CD8+ T cell properties, with high CD8+ T cell recruitment rate and PD1 blockade implemented. *dp*, probability of CD8+ T cell death; *kp*, probability of CD8+ T cell-mediated cancer cell killing. Note the *y*-axes limits are set to enable direct comparison with Figures 1, 2, and S2.

**Figure 8.**
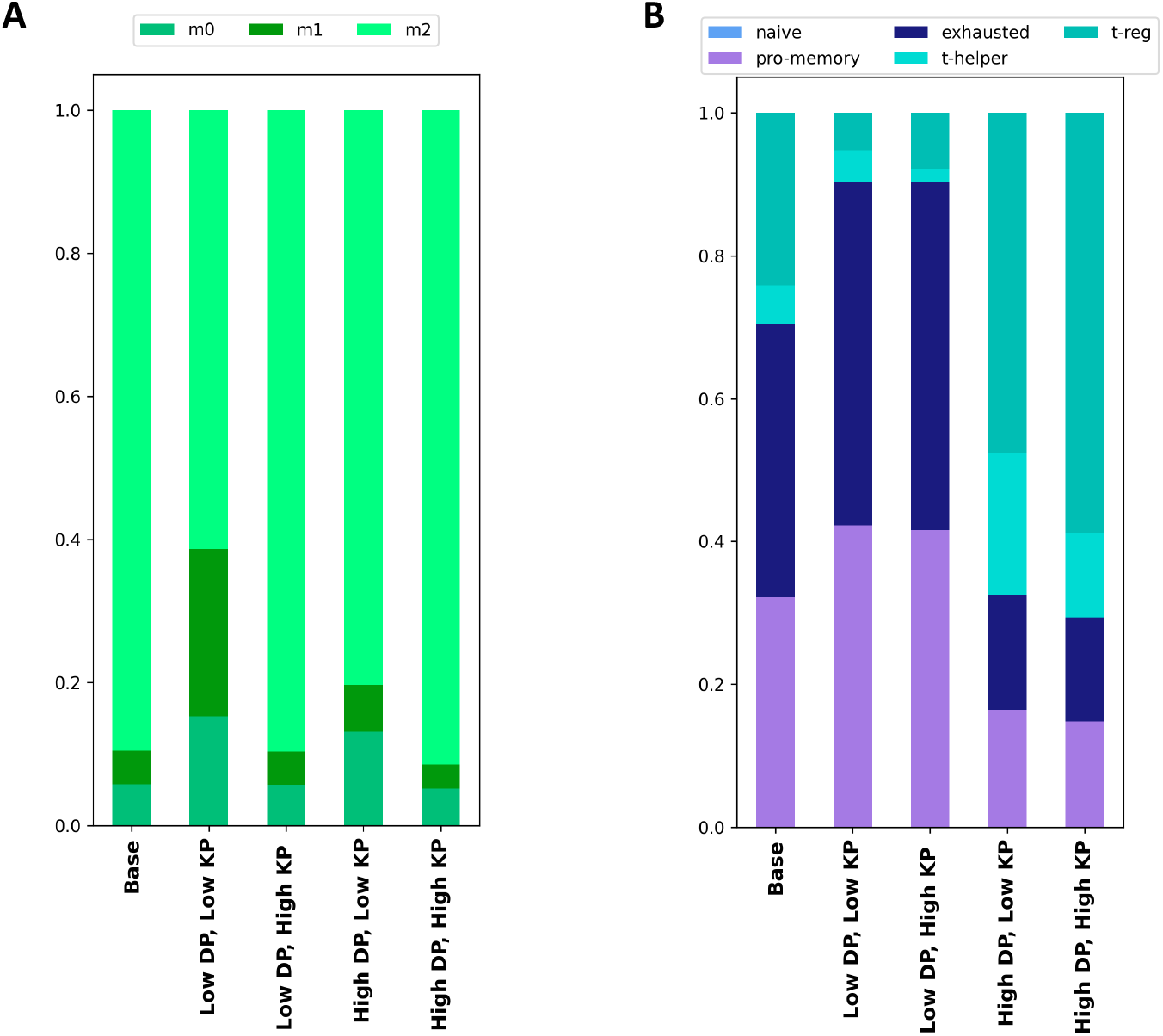
Relative proportions of immune cells at the end of the model simulation for various CD8+ T cell properties, with high CD8+ T cell recruitment rate and PD1 blockade implemented. **A**, Proportions of macrophages. **B**, Proportions of T cells. *dp*, probability of CD8+ T cell death; *kp*, probability of CD8+ T cell-mediated cancer cell killing.

In contrast, PD1 blockade consistently increases the fraction of pro-memory CD8+ T cells (compare **Figures 4B and 8B**). The condition of having a low probability of CD8+ T cell death and high probability of T cell-mediated killing is an exemplary case. Here, the percentages of T cells in the pro-memory state at the end of the model simulation are 1% and 34%, respectively, for the WT and PD1 blockade cases. Visual inspection of the spatial layouts also demonstrates the effect of PD1 blockade (**Figure S4**). We note that for a high probability of death CD8+ T cell death, PD1 blockade is not able to reduce the number of cancer cells (top row of **Figure 7D, E**), despite there being more pro-memory CD8+ T cells. In summary, a combination of high CD8+ T cell recruitment and low CD8+ T cell death is needed for PD1 blockade to effectively control the size of the cancer cell population.

## 3 DISCUSSION

We have presented an agent-based modeling framework that predicts the spatio-temporal behavior of the tumor-immune ecosystem comprised of cancer cells and distinct subsets of macrophages and T cells. The model highlights that CD8+ T cell recruitment, death and cancer cell-mediated killing can strongly influence tumor growth. In addition to predicting how the number of cancer cells evolves with time, our results consider the absolute number and relative proportions of macrophage and T cell subsets, representing the immune component of the tumor. Experimental and clinical studies demonstrate the efficacy of targeting the immune checkpoint inhibitor, PD1, to inhibit tumor growth.^22^ Therefore, we applied the model to predict the effects of altering gene regulation within CD8+ T cells to inhibit activation of the PD1 gene. This strategy is predicted to permit better control of the cancer cell population, in agreement with published experimental studies. Given the mechanistic basis of our model, we can explain the efficacy of PD1 blockade, finding that it drastically shifts CD8+ T cells from an exhausted state to a pro-memory state. Importantly, the model shows that the efficacy of this PD1 blockade in controlling the number of cancer cells depends on CD8+ T cell properties. This result also aligns with experimental studies, where engineering CD8+ T cell characteristics is used to enhance the effects of immune checkpoint blockade.^24^

A defining aspect of our work is the use of a Boolean model to inform the state transitions of CD8+ T cells. Specifically, we incorporate a Boolean representation of the GRN that mediates progression of CD8+ T cells from a naïve state to a terminally differentiated exhausted or pro-memory state.^20^ The T cell’s state influences its phenotype (recruitment rate and probability of cell death) and cell-cell interactions (probability of promoting cancer cell killing). Thus, integration of the Boolean model allows the behaviors of CD8+ T cells in the ABM to be directly determined by the intracellular gene regulation rather than taking a purely rule-based approach. Furthermore, this integrated approach enables the investigation of targeted genetic strategies to alter a T cell’s state. We have previously incorporated an intracellular network within an ABM to study the population-level effects of therapies that target macrophage intracellular signaling.^17^ Similarly, some ABMs consider intracellular models to inform the agent’s behaviors.[refs] However, most ABMs use phenomenological rules to determine cell state changes and phenotypes. Thus, having a mechanistic basis for the CD8+ T cell behavior helps advance the field of agent-based modeling.

We acknowledge some areas for future improvements of this work. Firstly, our model includes three classes of macrophages and five types of T cells. This is a vast simplification of the tumor-immune ecosystem. Thus, we can expand the model to include other immune subsets. For example, recent work reveals that interactions between T cells and dendritic cells influence T cell state changes and tumor growth.^25,26^ Related to this, we include differentiation of macrophages into just two broad classes (M1 and M2, anti- and pro-tumor, respectively); however, future work can consider a range of macrophage states and behaviors. Aligning the time steps simulated by the Boolean model is a second limitation of our work. Based on published experimental findings, we model that CD8+ T cells become exhausted within 9 hours of stimulation by a cancer cell based on the findings of X and Y.^27^ An issue with Boolean modeling is the use of pseudo-time, where the discrete time steps do not directly match real time. If the time required to reach an attractor state is known, one must assume what the interval between Boolean pseudo-time step corresponds to. For simplicity, we assume linear pseudo-time steps and that the Boolean model time steps correspond to a time interval of one hour. While the current work focuses on phenotypic features of CD8+ T cells, the effects of the time intervals within the Boolean model and aligning the timing between Boolean and ABM can be explored in future work. Thirdly, we note that altering the GRN that mediates CD8+ T cell state transitions may not happen with 100% efficiency in a heterogeneous population of cells. We can account for this by sampling from 10^4^ simulated trajectories that are a mixture of WT and PD1 blockade. For example, assuming an alteration to the GRN within a population of cells is 80% effective, we would sample from a set of trajectories comprised of 2×10^3^ WT and 8×10^3^ PD1 blockade Boolean simulations. Finally, this work focuses on development of the integrated Boolean-ABM framework and explores how CD8+ T cell properties affect tumor growth. We have not performed a full analysis of the effect of varying all model parameters and have not rigorously calibrated the model to tumor images from experimental models such as organoids or *in vivo* mouse tumors. Future work will leverage our recently developed machine learning-based approach to calibrate ABMs to such imaging data.^19^

## 4 CONCLUSIONS

The Taking an ecological view of a biological system by considering its spatial organization provides a framework for understanding the evolution of the system and the factors that control system behavior.^28^ Similarly, in the context of cancer, by considering the composition and spatial structure of tumors, it may be possible to better understand the factors that drive tumor progression and response to treatment.^7^ We have demonstrated that biophysical agent-based modeling can produce biological insights into the tumor-immune ecosystem. Furthermore, accounting for the evolution of T cell states as determined by an intracellular model of CD8+ T cell gene activation simulates experimentally observed behaviors. Overall, our work provides a solid foundation for future studies to explore the role of cell-cell interactions in the TME and how they influence the response to treatment.

## 5 METHODS

### Model Description

*Model Overview*. We develop an ABM to simulate the actions and interactions between cellular species in the TME. Specifically, we are able to capture the emergent outcomes associated with the interactions between cancer cells, cytotoxic CD8+ T cells (also referred to as CTLs), helper CD4+ T cells, and macrophages classed into three phenotypes (M0, M1 and M2). The cell-cell interactions are depicted in **Figure 9**. This model provides a framework to develop sophisticated simulations of the TME and capture complex biophysical phenomena in simplified forms. We describe the model features below.

**Figure 9.**
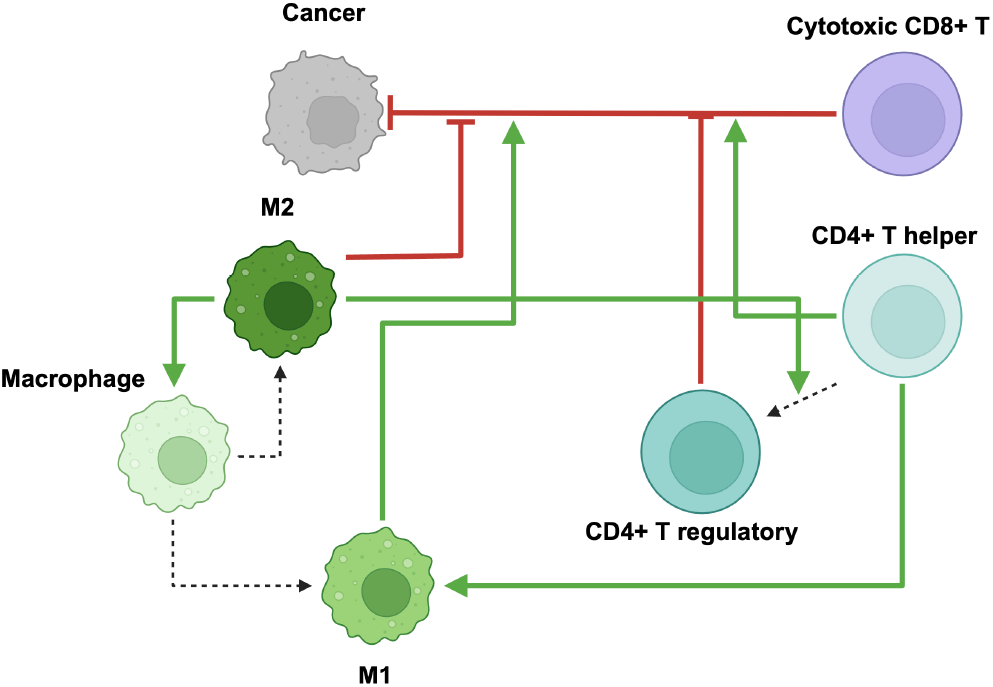
Schematic of cell-cell interactions included in the ABM.

*Cell Forces*. Cells are modeled using a center-based approach that considers each cell as a point and a radius; these two features are used to calculate physical forces between cells.^18,29^ This approach provides greater level of biological realism compared to grid-based approaches like the cellular-automata method. At the same time, the center-based approach is not as computationally expensive as frameworks that account for more detailed cell shapes such as the vertex model.^29^ Intercellular forces are calculated via the following equation.

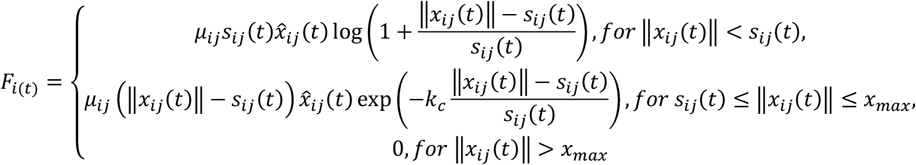

In this equation, *μ*_*ij*_ is the spring constant, *x*_*ij*_(*t*) is the vector between cells *i* and *j* at time *t*, 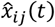 is the unit vector, *k*_*c*_ is the decay of the attractive force, *s*_*ij*_(*t*) is the sum of the radii of cells *i* and *j*, and *x*_*max*_ is the maximum interaction distance. For all cells, we assume the same *μ* and *k*_*c*_. This equation is solved numerically for each cell, as displayed in the following equation.

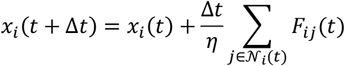

Here, *η* is the drag coefficient and *𝒩*_*i*_ is the set of cells within *x*_*max*_ of cell *i*. To preserve accuracy, this equation must be solved at a small timestep (Δ*t*), which we set as 0.005 hours. The parameters *x*_*max*_, *η, μ*, and *k*_*c*_ are taken from literature.

*Proliferation and cell death*. To simplify proliferation and cell death, these processes are modeled as probabilities of occurring at each time point. Cell proliferation is also influenced by the physical presence of other cells, assuming that cell proliferation is inhibited by excess physical forces.^18^ If the total overlap with other cells is above a threshold, a cell’s proliferation is inhibited until this overlap decreases. When a cell proliferates, the daughter cell is placed the distance of the cell radius away from the mother cell at a randomly selected angle. When a cell dies, it is removed from the simulation. Only cancer cells and CD8+ T cells can undergo proliferation.

*Immune cell recruitment*. Immune cells are recruited to the environment at a rate proportional to the number of cancer cells. They are recruited at a random angle around the tumor center, under the assumption that the area surrounding the tumor is well vascularized. When recruited, these cells are placed a random distance away from the tumor edge, sampled from a uniform distribution.

*Immune cell migration*. In the model, immune cells migrate towards tumor cells, mimicking chemotaxis due to tumor-secreted chemokines.^30–32^ Cells migrate with a specified amount of bias, which is modeled as the probability at each migration step of moving in the direction of the tumor center versus a random direction. This represents the impact of the extracellular matrix surrounding the tumor.^33–37^ However, there are many barriers to immune cell infiltration into the tumor, leading to three general tumor-immune states: ignored (immune cells cannot detect the tumor), inflamed (immune cells can infiltrate the tumor) and excluded (immune cells are restricted to the outside of the tumor).^38^ Some of these are due to interactions between immune cells, while others are due to physical barriers of the tumor. To capture this phenomenon in a simplistic manner with as few additional parameters as possible, we assume that both migration speed and bias decrease upon entry into the tumor. By adjusting these decreases, this model can capture different phenotypic tumor behaviors. Migration for all immune cell types is modeled in the same way; however, each cell type has its own bias and speed parameters.

*Diffusion*. Within the TME, cells can communicate via diffusible factors, such as cytokines.^32^ These are often modeled using PDEs, with cells acting as point sources/sinks.^10,13,39^ PDEs, however, require a great amount of computational time to solve, especially as environment size and the number of factors increases. Additionally, cells secrete several different cytokines, many with overlapping effects. This complexity makes it difficult to model the explicit biological effects of all the cytokines. Often, the effects of these factors are modeled either as a probability or a threshold value, instead of explicit modeling of downstream intracellular signaling. Because of this, researchers lump different cytokines into generic factors,^39^ or replace diffusible factors with a distance threshold, where an effect occurs if one cell is within a specified distance from another cell.^40^

Here, we implemented and extended the latter approach to better account for the gradient nature of diffusion and the impact of multiple secreting cells. We refer to this as the “cell influence,” where the closer cell *i* is to cytokine-secreting cell *j*, the greater the effect of cell *j*’s on cell *i*. This is modeled by an exponential decay displayed in the following equations:

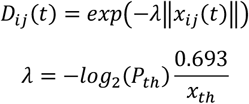

The use of an exponential decay is a reasonable approximation of the effects of diffusible factors, as the response to a cytokine-secreting cell has been experimentally found to resemble an exponential decay.^41,42^ In these equations, ‖*x*_*ij*_(*t*) ‖ is the distance between cell centers, *x*_*th*_ is the soft threshold for the maximum influence distance that can be thought of as the diffusion limit, and *P*_*th*_ is the probability of an effect occurring at distance *x*_*th*_. By setting *P*_*th*_ and *x*_*th*_, we can calculate *λ*. The total influence on cell *i* is calculated via

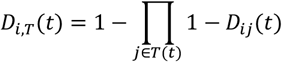

In this equation, *D*_*i,T*_(*t*) is the total influence, from 0 to 1, on cell *i* from all cells of cell type *T*. Thus, each cell records a separate influence for each cell type in the simulation. These influences are used to determine downstream effects. These downstream effects are probabilities, which are then multiplied by the relevant influence. Thus, the cell influence acts as a scaling factor for the probabilities of effects occurring. If two cell types can lead to the same effect, their influences are combined as

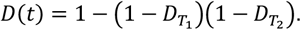

This approach eliminates numerically solving PDEs and means that the cloud of diffusible factors follows each cell as it moves through the simulation environment. We followed the assumption that diffusion is resolved on a much faster timescale than cellular processes.

*Cancer Cells*. In this model, cancer cells have three main actions: proliferation, death, and expression of the PD1 ligand (PD-L1). Proliferation and death occur as described above. Cancer cells gain PD-L1 at a probability proportional to the influence from CD4+ helper and CD8+ cells (via T cell-secreted IFN-γ, which promotes PD-L1 expression).^10,43^ PD-L1 expression is represented as a probability of suppressing CD8+ cells upon direct contact.^10^ After proliferation, the daughter cells inherit the PD-L1 expression of the mother.

*Macrophages*. Classical *in vitro* characterization of macrophages leads to their classification into what have been called M1 pro-tumor and M2 anti-tumor macrophages. While it is now known that macrophages exist on a continuum, whereby their functions vary, and they can exhibit a range of phenotypes and functions,^44,45^ we retain these broad M1 and M2 groupings for the purposes of exploring the effects of anti- and pro-tumor macrophages.

Macrophages have a certain size^46^ and enter the simulation in a naïve (M0) state and differentiate into either M1 (mimicking the response to IFN-γ secretion by CD4+ helper and CD8+ T cells) or M2 (mimicking the response to IL-4 and IL-10 secreted by cancer cells and Treg cells).^47^ Additionally, because macrophage state is plastic and influenced by environmental conditions, macrophages have a probability of re-differentiating at each timestep.^48^ Following other modeling efforts, macrophages have a probability of remaining in the naïve (static value) or differentiating into M1 or M2 (influenced by environmental conditions), which are then scaled to sum to 1.^13^ This is captured by the following equations, which utilize the cell influence described above.

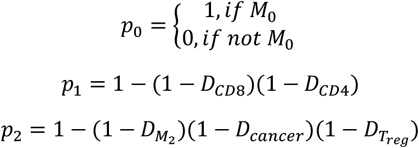

These values are then divided by their sum to get the probability of differentiating into each state.

M1 cells increase the killing capability of CD8+ T cells (via cytokines such as IL-12)^47,49^ and are able to kill nearby cancer cells (via nitric oxide secretion). M2 cells reduce the killing capability of CD8+ T cells and proliferation rate of CD8+ T cells (via IL-10 and TGF-β), promote the differentiation of CD4+ T cells into the regulatory state (via IL-10 and TGF-β), and promote M0 differentiation into the M2 state (via TGF-β).^47,50,51^ We model all of these effects via the cell influence function described above. M2 cells also express PD-L1, represented as a probability of suppressing a CD8+ cell upon direct contact. *CD4+ T cells*. While CD4+ T cells can take on a variety of states, for simplicity, we only model two: helper cells and regulatory cells. In this model, CD4+ cells enter the simulation in the helper state and can be converted into regulatory cells based on influence from M2 macrophages and cancer cells.^47,50–52^ The probability of differentiation is proportional to the combined influence of M2 macrophages and cancer cells, as shown in the following equations.

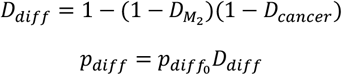

In the helper state, CD4+ cells promote the killing capability and proliferation of CD8+ T cells (via IL-2)^53–55^ and promote M0 differentiation to the M1 state (via IFN-γ).^56^ As regulatory cells, they express CTLA-4 (which has the same function as PD-L1),^57^ decrease the killing and proliferative capabilities of CD8+ T cells (via acting as an IL-2 sink),^53^ and promote M0 differentiation into the M2 state (via IL-10).^52^

*CD8+ T Cells*. CD8+ T cells serve the main function of killing cancer cells. In the TME, we consider a CD8+ T cell to be in a naïve, exhausted, or pro-memory state. CD8+ T cells can undergo cell death, kill a neighboring cancer cell, and migrate^58^. The probability of each of these two behaviors is determined by the CD8+ T cell state. Specifically, we set the model parameter for each of these cell behaviors relative to a baseline set of behaviors, further detailed in the “*Integrating CD8+ T cell state decision into ABM*” below. Finally, When a CD8 cell comes into contact with a PD-L1 expressing cell, the CD8 cell becomes suppressed based on the probability of PD-L1.^10^

*Defining the Simulation Loop*

The simulation loop proceeds as follows (**Figure 10**):

1. Immune cells are recruited to the environment.
2. Cell neighborhoods are determined.
3. Influence from cell-cell interactions due to proximity (mimicking the effects of diffusible factors) is calculated, direct contact effects (CD8+ T cell-mediated cancer cell killing, CD4+ differentiation) are accounted for, CD8+ T cell states are updated based on intracellular Boolean model.
4. Forces between cells are solved at a small timestep.
5. Species proliferation as well as spontaneous cell death are modeled.

**Figure 10.**
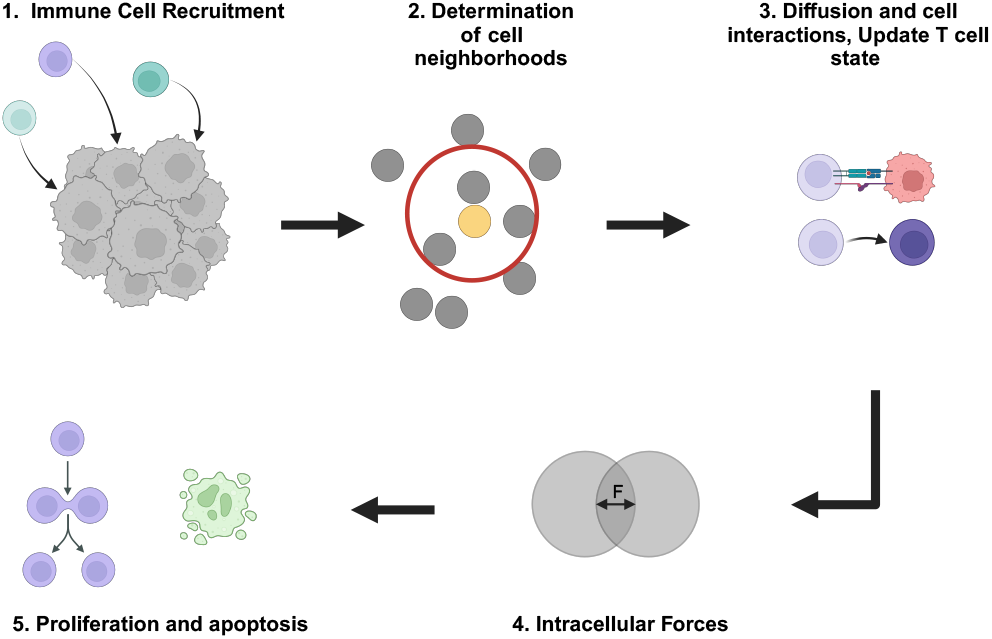
Illustration of steps followed in the ABM simulation loop.

### Predicting CD8+ T cell states

CD8+ T cell state decisions were determined based on the output of our previously published Boolean model of the GRN that defines CD8+ T cell states.^20^ This model predicts the expression profile of CD8+ T cell genes over pseudo-time. Although CD8+ T cells occupy a continuum of phenotypes, we broadly class the cell’s state into one of three main phenotypes: naïve (N), pro-memory (M), and exhausted (E), based on the output of the GRN predicted by the Boolean model. The model logs the on/off expression of 24 genes, where each gene is classified as promoting a proliferative pro-memory (PP) state or an effector exhausted (EE) state. The Boolean model predicts the state of each CD8+ T cell at every time step based on the relative expression of PP and EE genes. Specifically, if more PP genes are *on* compared to EE, the phenotype is assumed to be M; the phenotype is assumed to be E if there are more EE genes *on* than PP. The first three pseudo-time steps are assumed to be in the N state; following stimulation of the T cell receptor, the predicted state in subsequent time steps is based on the expression of PP and EE genes.

For a single simulation, the output from a Boolean model simulation is given by an *n* × *k* matrix where *n* represents the number of genes (24 in total) and *k* represents the number of time steps. The (*n, k*) entry in the matrix is the CD8+ T cell state (N, P, or E), given by the rules defined above. Thus, moving down the rows of this matrix represents the “trajectory” that a cell takes in its evolution from a naïve state to the terminally differentiated state. Once a cell reaches its terminal state, it stays in that state for the remaining *n* time steps. Our prior work demonstrated that the Boolean model captures both linear differentiation of a T cell from M to E, and circular differentiation where the cell oscillates between M and E before reaching the final state.^59,60^

Ten thousand simulations of the Boolean model were performed, representing a population of 10^4^ distinct CD8+ T cells. We first consider the baseline or wildtype (WT). We also considered the case where the CD8+ T cell GRN encoded by the Boolean model reflects a PD1 blockade. Literature evidence shows that nuclear factor of activated T cells 1 (NFATC1) can promote antitumoral effector functions and memory CD8+ T cell differentiation through regulation of PD1 expression upon T cell activation.^61,62^ Therefore, as described in our prior work,^20^ we implemented indirect inhibition of PD1 via NFATC1. In the WT model, NFATC1 can activate PD1 (NFATC1 → PD1), and here, we removed NFATC1 from the model, inhibiting PD1 activation. With this alteration of the GRN, 10^4^ additional simulations of the Boolean model were performed to produce a population of 10^4^ distinct CD8+ T cells in which PD1 is inhibited.

### Integrating CD8+ T cell state decision into ABM

*Updating ABM based on state predicted by Boolean model*. The predicted CD8+ T cell state given by the Boolean model transitions through pseudo-time. It is straightforward to initialize every CD8+ T cell that is recruited to the tumor by sampling from the 10^4^ trajectories produced by the Boolean model and assigning the selected trajectory to the CD8+ T cell. However, we must assign a time interval for each pseudo-time step to align the transition of CD8+ T cells states with the timescale of the ABM. Experimental evidence indicates that CD8+ T cells become exhausted within six to 12 hours of stimulation by a tumor cell.^27^ We reflected this finding by updating the CD8+ T cell state in the ABM every 9 hours, assuming that each pseudo-time step of the Boolean model corresponds to one hour. Thus, the predicted CD8+ T cell state given by the Boolean model at every ninth time step is passed into the ABM. In the cases where the ABM time point is longer than the trajectory predicted by the Boolean (i.e., the CD8+ T cell has reached its terminally differentiated state before the current ABM simulation timepoint), we assume the T cell remains in its final state (the last value in its trajectory predicted by the Boolean model) for the rest of the simulation until cell death. For PD1 inhibition, we instead initialize and update the states of CD8+ T cells using the Boolean simulations corresponding to the altered GRN in which PD1 activation by NFATC1 is removed.

*Assigning cell behavior based on state predicted by Boolean model*. The “*Model Description*” section details that CD8+ T cells can die and kill cancer cells. The probability of each of these behaviors is determined by the state predicted by the Boolean model, relative to a base probability. CD8+ T cells in the N state are assumed to show no deviance from the base probabilities. In comparison, cells in the M state are half as likely to die and twice as likely to trigger cancer cell death, relative to baseline. Finally, CD8+ T cells in the E state maintain the base probability of death; however, their ability to promote cancer cell killing is reduced by an order of magnitude, relative to baseline. By varying cell death and cytotoxicity, we capture CD8+ T cell state-specific behaviors.

### Model Implementation

This model is built in C++ using CMake for build processes. Scripts that develop the parameter grid as well as plotting scripts were implemented in Python. Two builds are presented, one developed for Apple Silicon and the other for a Unix-based computing cluster. The code is available at: https://github.com/FinleyLabUSC/Boolean-TME-ABM.

### Model Simulation

The parameter values used in the model are given in **Table S2**. We perform simulations to explore the effects of CD8+ T cell properties: recruitment rate (*k*_recr_), probability of cell death (*DP*), and cancer cell killing probability (*KP*). We further investigate increased T cell recruitment (5-fold higher than the baseline value); varying the probability of cell death: 5-fold increase and reduction in *DP* (high *DP* and low *DP*, respectively); and varying the probability of cancer cell killing: 5-fold increase and reduction in *KP* (high *KP* and low *KP*, respectively). To account for variability in the predictions due to the probabilistic nature of the agent-based modeling, we performed ten replicates for each parameter combination considered.

## ACKNOWLEDGEMENTS

We acknowledge the Finley research group for critical evaluation of this manuscript. This work was supported by the USC Center for Computational Modeling of Cancer.

## AUTHOR DECLARATIONS

The authors have no conflicts to disclose.

## DATA AVAILABILITY STATEMENT

The data that supports the findings of this study are available within the article and openly available on GitHub at https://github.com/FinleyLabUSC/Boolean-TME-ABM.

## SUPPLEMENTARY MATERIALS

**File S1:** Movie 1 – Representative time course of spatial layout over time for the baseline probabilities of CD8+ T cell death and cancer cell killing for base CD8+ T cell recruitment. Provided as a .mp4 file

**File S2:** Movie 2 – Representative time course of spatial layout over time for the baseline probabilities of CD8+ T cell death and cancer cell killing for high CD8+ T cell recruitment. Provided as a .mp4 file

**File S3:** Supplementary figures; provided as a .pdf file.

**File S4:** Table S1. Number and proportion of cell types, and mean infiltration distance for immune cells at the end of model simulations.

**File S5:** Table S2. Model parameters.

