## Supplementary material for "Integrating mechanism-based T cell phenotypes into a model of tumor-immune cell interactions": File S3

### SUPPLEMENTARY FIGURES

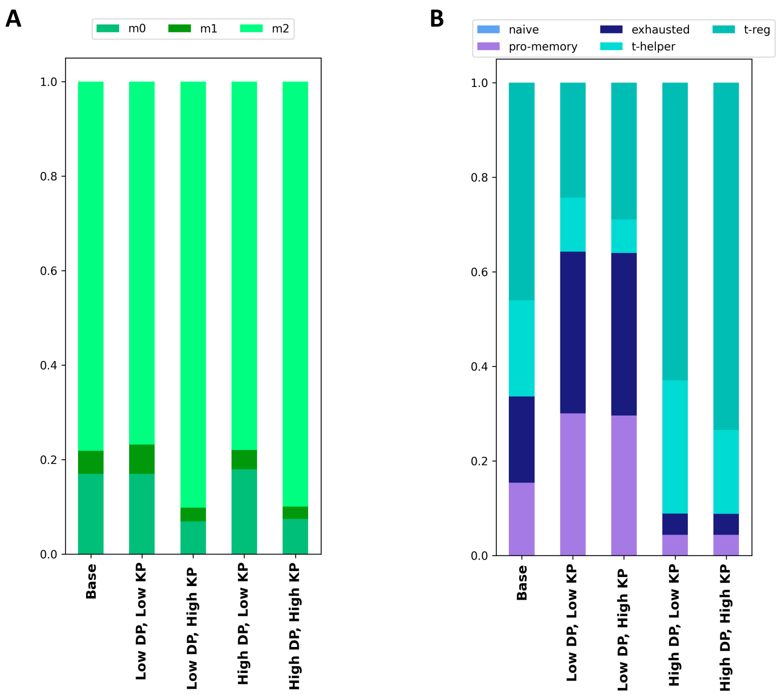


**Figure S1:** Relative proportions of immune cell populations at the end of the model simulation for various CD8+ T cell properties, for base CD8+ T cell recruitment rate with PD1 blockade. **A**, Proportions of macrophages. **B**, Proportions of T cells. *dp*, probability of CD8+ T cell death; *kp*, probability of CD8+ T cell-mediated cancer cell killing.

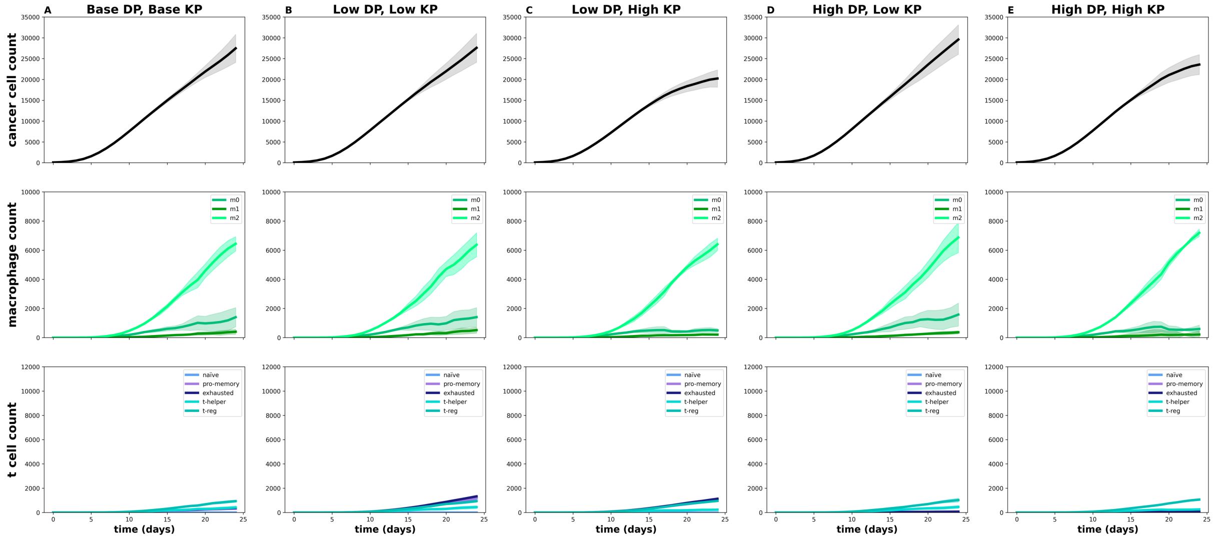


**Figure S2:** Time courses of cell counts for various CD8+ T cell properties, for base CD8+ T cell recruitment rate with PD1 blockade. *dp*, probability of CD8+ T cell death; *kp*, probability of CD8+ T cell-mediated cancer cell killing.

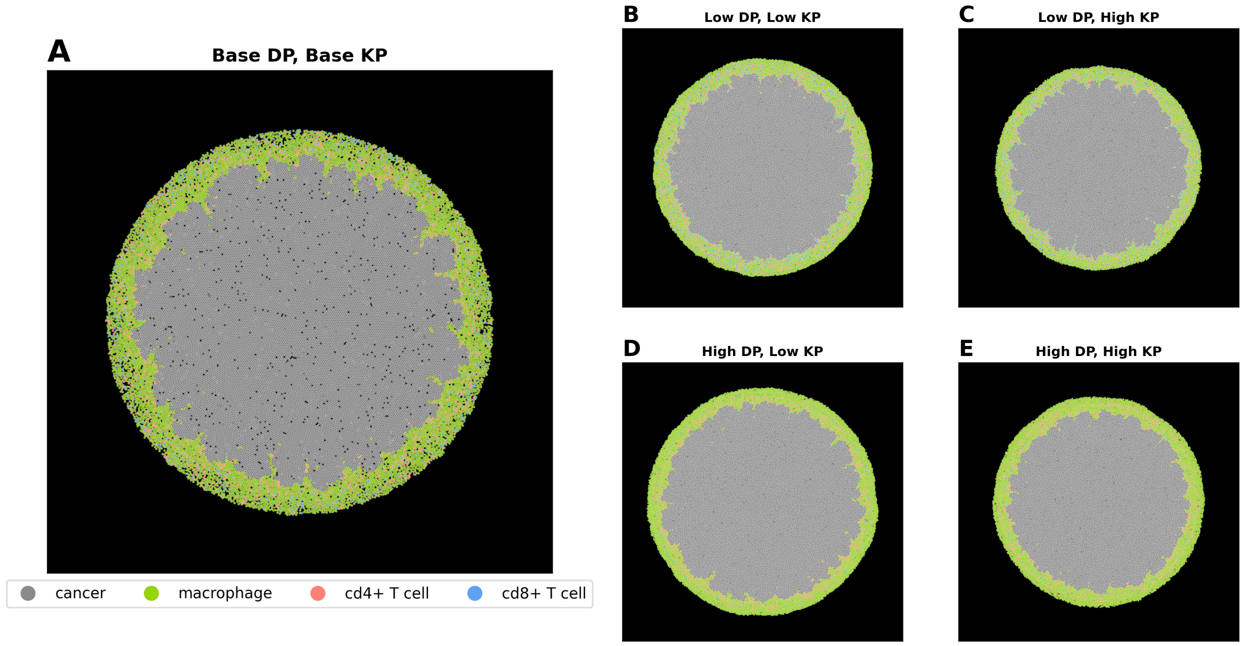


**Figure S3:** Representative tumor spatial layouts for various CD8+ T cell properties at the end of the simulation, for low CD8+ T cell recruitment rate with PD1 blockade. *dp*, probability of CD8+ T cell death; *kp*, probability of CD8+ T cell-mediated cancer cell killing.

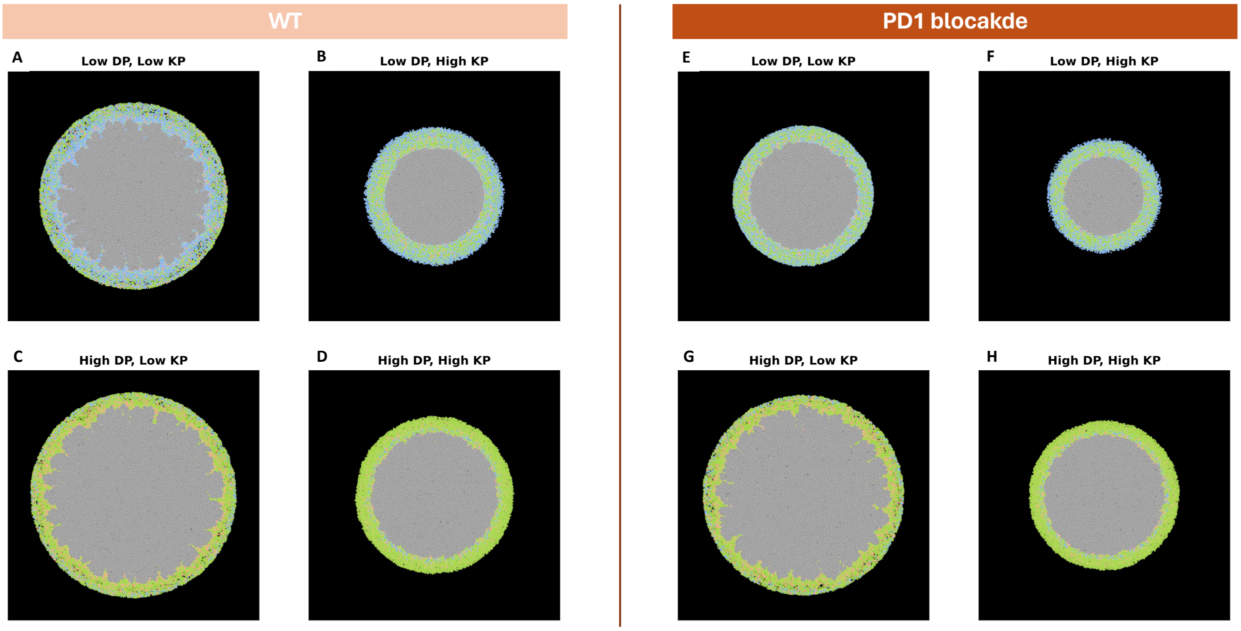


**Figure S4:** Representative tumor spatial layouts for various CD8+ T cell properties at the end of the simulation for high CD8+ T cell recruitment. **Left**, WT baseline model. **Right**, PD1. *dp*, probability of CD8+ T cell death; *kp*, probability of CD8+ T cell-mediated cancer cell killing.
